# Minus one frameshifted puromycin N-acetyltransferase is targeted to the nucleolus: sporadic but reproducible chimera formation by a transfected *Lc3b* construct

**DOI:** 10.1101/2022.02.17.480764

**Authors:** Arvind A. Thekkinghat, Pundi N. Rangarajan

## Abstract

Map1Lc3b is a protein that has pivotal functions in cellular autophagy. At least three groups in the past decade have reported its presence in the nucleoli of cells, but its functions in that organelle remain unknown. We isolated a few clonal populations of cells stably expressing V5-tagged mouse Lc3b highly enriched in the nucleoli, but the frequency of occurrence of such clones was strikingly low. The phenomenon was readily reproducible, though the protein in the nucleolus puzzlingly had varying molecular masses in different clones but consistently displayed a very strong interaction with the mitochondrial protein C1qbp, which has well-documented functions in the nucleolus. We investigated further and discovered that, in at least one of the clones, *Lc3b* had formed a chimera with the puromycin resistance gene in the plasmid, plausibly by illegitimate recombination during or after integration of the construct into the cellular genomic DNA. The -1 shifted reading frame of puromycin N-acetyltransferase (*pac*) can encode a protein that is equally long as the one encoded by the complete *pac* ORF, but is targeted to the nucleoli due to a drastic shift in the isoelectric point (pI). Notably, this set of events again brings into focus the low threshold often reported for recombination events to occur in eukaryotic cells, the multiple factors influencing them, and calls for increased vigilance in experiments involving DNA transfection and gene targeting.

## Introduction

MAP1LC3B is an important protein in mammalian cells and is widely recognised as a crucial component of autophagosomes where it decorates the membranes, driving the process of autophagic degradation by interacting with a variety of autophagic cargo receptors and other proteins that modulate the specificity and kinetics of the different types of autophagy (Kabeya, 2004; Birgisdottir et al., 2013, Khaminets et al., 2016). MAP1LC3B (often referred to as just LC3B) and its family members MAP1LC3A, MAP1LC3C, GABARAP, GABARAPL1, and GABARAPL2 are ubiquitin-like proteins that are orthologs of yeast Atg8. Lipidation of these proteins involving covalent attachment to phosphatidylethanolamine enables them to become membrane-bound during the process of autophagy (Kabeya et al., 2004), serving as a marker for the process. At the same time there are indications that these proteins might have other functions within the cell, in addition to their much vaunted roles in autophagy. In 2015, Koukourakis et al. performed studies on multiple human cell lines and reported that the three proteins of the LC3 family – LC3A, LC3B, and LC3C – had distinct and partially non-overlapping subcellular distribution patterns inside the cell, indicating a possibility of non-identical kinetics and distinct biological roles for each member (Koukourakis et al., 2015). A notable finding reported by the authors in this paper was that LC3B, unlike the other two members, was present in the nucleoli in addition to being distributed in the cytoplasm, though the percentage of cells with nucleolar localization varied among cell lines. Around the same time, Kraft et al., in a study of the nucleo-cytoplasmic trafficking and dynamics of LC3B and its association with large complexes in the cell, found that exogenously expressed Venus-LC3B was weakly enriched in the nucleolus (Kraft et al., 2016). They commented that the relatively weak association of LC3B with the nucleolus might be a reason for it being overlooked in previous studies, and that it might interact with one or more protein/RNA components in that organelle. It was also noted that the mutation of a triple arginine motif R68-70 in LC3B, which had previously been shown to bind an AU-rich region in the 3’-UTR of fibronectin mRNA (Zhou et al., 1997), completely disrupted nucleolar localization of LC3B. In 2019, Shim et al. reported that LC3 localizes to the nucleoli of cells of the trabecular meshwork of the anterior chamber of the eye, in response to stretching of the cells by cyclic mechanical stress (Shim et al., 2019).

In 2012, we had isolated a few clonal populations of cell lines transfected with mouse *Lc3b* to stably express the gene, in which it appeared that Lc3b was unusually highly enriched in the nucleoli compared with other regions in the cell. Later we investigated what might be causing this unusual expression pattern, and how Lc3b in these cell lines might differ from the norm. To our surprise we discovered that, unlike the observations of Koukourakis et al. and Kraft et al. which likely reflect true nucleolar localization of Lc3b, our observations were the result of a very specific, highly reproducible, and might we say rather spectacular, artefact in which the plasmid used for transfection, the epitope tag, and a chimera comprised of two partial reading frames all contributed to create a deceptive illusion of Lc3b localizing to the nucleolus.

## Results

### 3.1 Stably transfected V5-tagged *Lc3b* yields clonal populations with Lc3b highly enriched in the nucleoli, at a very low incidence

Mouse *Lc3b* was tagged with a V5 tag at its N-terminus, transfected into B16F10 melanoma cells and selected for stable expression with 2 μg ml^-1^ puromycin. The normal pattern of localization of V5-Lc3b in B16F10 cells is essentially diffuse distribution in the cytoplasm and nucleus, with a moderate number of granular ALIS (Aggresome-Like Induced Structures)-like structures in the cytoplasm that persist in normal growth conditions **(Fig. 1A)**. But, in very few cells, V5-Lc3b showed a pattern of strong enrichment in the nucleoli **(Fig. 1B)**. We estimate that this low incidence is somewhere between 1-in-40 to 1-in-100 transfected cells (empirical estimation). Considering the low frequency of occurrence, to narrow this down to a more precise number would need isolation and expansion of a large sample size of clonal populations of Lc3b-expressing cells over multiple rounds of experiments, followed by immunofluorescence (IF) screening for nucleolar signal. Transiently lipofecting *V5-Lc3b* into B16F10 cells on a cover slip, followed by IF staining with anti-V5 antibody, reveals the odd transfected cell with nucleolar Lc3b in a large field of cells **(Fig. 1C)**; isolating a clonal population of such cells necessitates dilution of a transfected culture and plating at low density, followed by puromycin selection over ten days, picking single clones, expanding them, and screening by immunofluorescence for nucleolar staining.

**Fig. 1:**
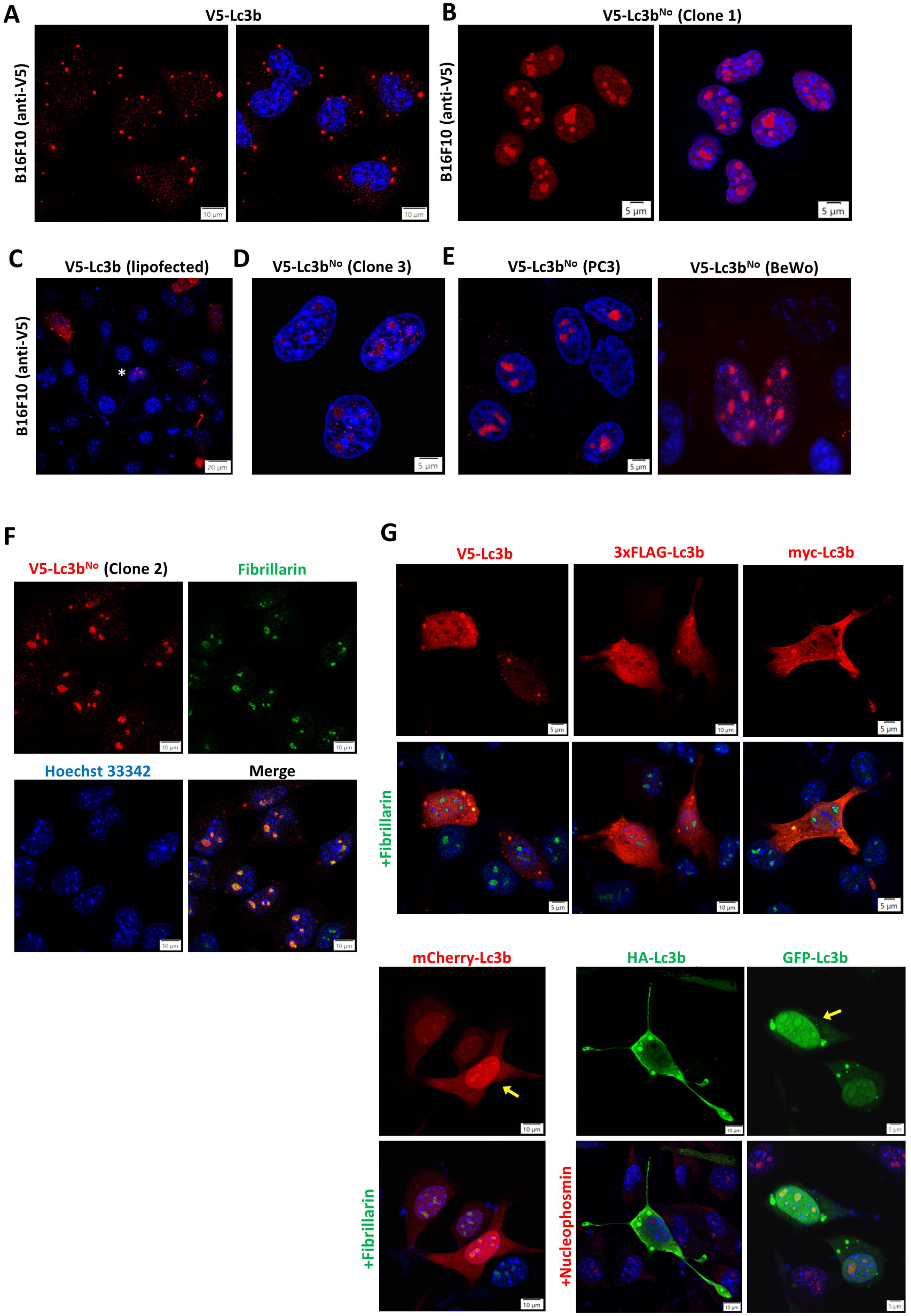
Subcellular distribution of V5-Lc3b in B16F10 melanoma cells. **A**. Typical distribution pattern of V5-Lc3b in B16F10 cells stably expressing the *V5-Lc3b* construct cloned in pBApo-EF1α Pur; red staining is with anti-V5 antibody. The panel on the right shows the nuclei stained with Hoechst 33342. **B**. Atypical distribution pattern (nucleolar) of V5-Lc3b in B16F10 cells stably expressing the *V5-Lc3b* construct (referred to as Lc3b^No^ for convenience; No = nucleolar). The panel on the right shows the nuclei stained with Hoechst 33342. This clone was expanded and passaged (‘clone 1’). **C**. B16F10 cells transiently transfected with the *V5-Lc3b* construct; note the odd cell (asterisked) in a large field exhibiting nucleolar staining. Almost all transfected cells show the typical Lc3b cytoplasmic/nuclear pattern. **D**. A clonal population of B16F10 expressing Lc3b^No^ (named clone 3) **E**. Clonal populations of BeWo and PC3 cells with a similar nucleolar staining pattern of stably expressed V5-Lc3b. The PC3 clone was isolated and passaged. **F**. Another clonal population of B16F10 cells (clone 2) with nucleolar Lc3b; nucleoli were stained by an anti-fibrillarin antibody. **G**. *Lc3b* constructs with different epitope tags and fusions transiently expressing the constructs after lipofection; Lc3b was detected with antibodies against the respective epitope tag, or by direct fluorescence from the fluorophore in the fusion protein. Nucleoli are stained by either anti-fibrillarin or anti-nucleophosmin, according to antibody compatibility with the contrasting fluorophore. Note the unstained nucleoli where Lc3b is tagged with small epitope tags, compared with GFP and mCherry fusions (yellow arrows).

To study this phenomenon, we isolated three clonal populations of B16F10 melanoma cells with V5-Lc3b predominantly localized in the nucleoli **(Fig. 1B, D, F)**. This phenomenon is not cell line-specific; we observed it at a similar low incidence when the construct was transfected into both human BeWo choriocarcinoma cells and PC3 prostate cancer cells **(Fig. 1E)**. Two clonal populations of PC3 cells expressing nucleolar Lc3b were isolated for further analysis. All clones were passaged for several weeks with the phenotype remaining intact. The protein co-localized well with nucleolar markers like fibrillarin **(Fig. 1F)** and nucleophosmin. For ease of communication, we term the Lc3b in these cells Lc3b^No^ (^No^ = nucleolar) to differentiate it from the typical (cytoplasmic-nuclear) Lc3b. Though it is clear that close to 99% of puromycin-selected V5-Lc3b clones exhibit normal cytoplasmic-nuclear staining that is typical to Lc3b, in order to rule out any incidental bias of the V5-tag towards nucleolar localization, we transiently transfected *Lc3b* tagged with different epitope tags/fusion proteins at the N-terminus into B16F10 cells and performed immunofluorescence microscopy **(Fig. 1G)**. Only GFP-Lc3b and mCherry-Lc3b showed any significant staining of Lc3b in the nucleoli; this was in addition to strong staining in other compartments. Lc3b tagged with other small tags (HA, myc, FLAG), including V5, was more or less excluded from the nucleoli, suggesting that the V5 tag does not, per se, influence nucleolar localization and that the phenomenon in question involves other unexplained mechanisms.

Kraft et al. had reported that the lipidation of LC3B at the C-terminus is unconnected with its nucleolar localization (Kraft et al., 2016); accordingly we found that a G120A mutation at the C-terminus of Lc3b (which prevents covalent attachment to lipid) had no bearing on the phenomenon described in this paper.

### 3.2 V5-Lc3b^No^ exhibits variation in molecular mass and interacts strongly with C1qbp/p32

The clones of B16F10 melanoma expressing V5-LC3b^No^ surprisingly showed proteins that displayed varying molecular masses when immunoprecipitated with anti-V5 agarose and in western blots of lysates probed with an anti-V5 antibody. Clone A (intense staining and with the protein most abundantly expressed) yielded a band running near ∼20 kDa, and clone B yielded a band near ∼37 kDa **(Fig. 2A)**. Clone C showed moderately good nucleolar fluorescence in microscopy, but for some reason proved to be difficult to detect in lysates through blotting/IP (likely suboptimal expression, coupled with inefficient extraction from the nucleolar compartment). The V5-Lc3b^No^ stably expressed in PC3 cells yielded a band just above 40 kDa (shown in Fig.2F). The V5-Lc3b^No^ bands were detected in western blots by both anti-V5 and anti-Lc3 antibodies **(Fig. 2B)**, suggesting that the antibodies were indeed detecting Lc3b.

**Fig. 2:**
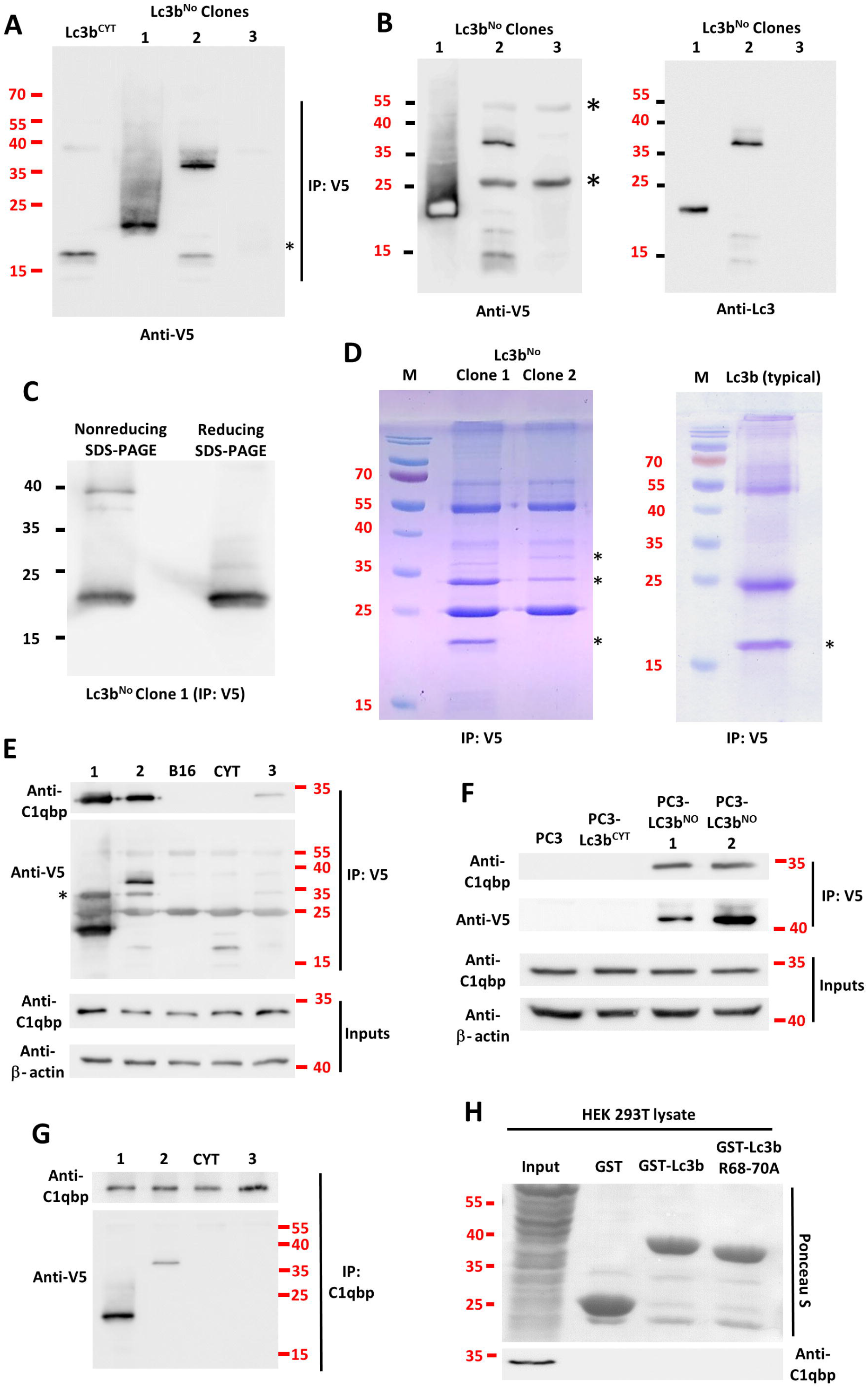
Biochemical characterization of V5-Lc3b^No^, and its interaction with C1qbp. **A**. Lc3b^No^ was immunoprecipitated (IP’ed) with anti-V5 agarose gel from cells of B16F10 Lc3b^No^ clones 1, 2 and 3 grown in 5x P100 plates. B16F10 cells stably expressing typical (cytoplasmic/nuclear) Lc3b was used as a control (labelled ‘Lc3b^CYT^’ in lane 1). The blot was probed with anti-V5. Asterisk indicates the position of very weakly stained band from clone 3. Only 25% of the IP’ed lysate was used for clone 1 because of extremely high expression levels. **B**. Lc3b^No^ was immunoprecipitated from the respective clones with anti-V5 agarose; the IP’ed protein was divided into two and electrophoresed on duplicate gels. One blot was probed with anti-V5, and the other with anti-Lc3b antibody. The overexposed band in lane 1 of the V5 blot is owing to very high levels of expression from clone 1. Asterisks indicate the positions of IgG heavy and light chain bands. **C**. Lc3b^No^ IP’ed from clone 1 was divided into two and run in two lanes on an SDS-PAGE – one non-reduced, and the other reduced with 2-mercaptoethanol. **D**. Lc3b^No^ was IP’ed from 8x P100 plates of clones 1 and 2, and the SDS-PAGE gel was stained with Coomassie Brilliant Blue. The three bands at positions with asterisks were excised and processed for mass spectrometry. The gel on the right shows the position of typical cytoplasmic-nuclear Lc3b (asterisked) IP’ed as a positive control. **E**. Lc3b was IP’ed with anti-V5 from 4x P100 plates of the respective B16F10 clones. Plain B16F10 cells were a negative control. The IP blot was first probed with anti-C1qbp and then anti-V5 without stripping (to retain weak signal from Lc3b^CYT^). Bands at 55 and 25 kDa indicate IgG chains. Asterisk indicates the position of the C1qbp bands. The inputs were run on a separate gel, and the blot stripped and re-probed for actin. **F**. Lc3b^No^ IP’ed with anti-V5 and probed for C1qbp, essentially as in E but from PC3 cells. The IP blot was stripped and re-probed with anti-V5. **G**. B16F10 clones IP’ed with anti-C1qbp and then probed for Lc3b^No^, essentially the reverse of E. **H**. Recombinant GST-Lc3b and GST-Lc3b^R68-70^ from *E. coli* were pulled down with glutathione-agarose and tested for interaction with C1qbp from HEK 293T cell lysates. GST was used as a control in lane 2. The experiments in this figure mostly have qualitative value only and do not gain additional value from quantitative analysis; however all IPs were performed with a minimum of three biological replicates and one representative blot has been shown.

The LC3b^No^ from clone A also yielded an extra band at a size approximately double its molecular mass when the samples were not reduced by 2-mercaptoethanol prior to SDS-PAGE, with a concomitant reduction in the intensity of the ∼20 kDa band **(Fig. 2C)**, suggesting that some of the protein might exist as a dimer.

When the protein was immunoprecipitated from these clones using anti-V5 agarose affinity gel, a protein of mass ∼32 kDa consistently co-precipitated with the V5-Lc3b^No^ bands from all the clones, but not with typical non-nucleolar V5-LC3b **(Fig. 2D-F)**. We observed that this interaction was so strong that it was well-conserved even when cell lysis was performed in denaturing conditions, i.e., by lysing the cells with a buffer containing 1% SDS, followed by subsequently diluting the lysate and neutralizing the SDS with an excess of Triton X-100 prior to immunoprecipitation (Bonifacino and Dell’Angelica, 1998). This step was deemed necessary because the V5-Lc3b^No^ in the nucleolus was not easily recovered in the soluble fractions of the cell lysates when using only non-ionic detergents in the lysis buffer, especially for clones other than clone A (which expressed a very high quantity of the tagged protein). Immunoprecipitating V5-Lc3b^No^ from 4-5 100 mm culture dishes yielded clear Coomassie Brilliant Blue-stainable bands of the IP’ed and co-IP’ed proteins on SDS-polyacrylamide gels **(Fig. 2D)**. Trypsin digestion followed by tandem mass spectrometry revealed the identity of the 32-kDa protein to be complement component 1, Q-subcomponent binding protein (C1qbp), also known as p32/HABP-1 (hyaluronan binding protein 1). The quantity of C1qbp co-immunoprecipitated was proportional to the quantity of V5-Lc3b^No^ expressed by the respective clones **(Fig. 2E)**, suggesting that it was specifically reacting with V5-Lc3b^No^.

Reverse-immunoprecipitation with an anti-C1qbp antibody readily co-immunoprecipitated V5-Lc3b^No^ from the Lc3b^No^ clones, further validating the interaction between the two proteins **(Fig. 2G)**. Though C1qbp is a mitochondrial protein, it has well-characterized functions in the nucleolus. C1qbp has been reported to participate in ribosome biogenesis in the nucleolus by regulating the binding of fibrillarin and Nop52 to preribosome particles (Yoshikawa et al., 2011). But C1qbp from cell lysates did not interact with recombinant glutathione S-transferase-tagged Lc3b (GST-Lc3b) in vitro **(Fig. 2H)**, confirming that the interaction is specifically with V5-Lc3b^No^.

### 3.3 Some inconsistencies and gaps cast doubt on the nucleolar localization of V5-Lc3b

The phenomenon of nucleolar localization of V5-Lc3b seemed genuine thus far, primarily because PCR from the cDNA or genomic DNA of the Lc3b^No^ clones with a V5-tag forward primer and the Lc3b reverse primer yielded an *Lc3b* band of the correct size in all the nucleolar clones **(Fig. 3A, B)**, indicating correct genomic integration and transcription from the construct encoding *V5-Lc3b*. Importantly, the proteins were detected in immunoblots by anti-Lc3 antibodies, in addition to the anti-V5 antibody (shown in the previous section). However, mass spectrometry of Lc3b^No^ bands immunoprecipitated by anti-V5 agarose consistently failed to unambiguously confirm the identity of the protein as Lc3b. None of the mass spectrometry outputs yielded Lc3b as the top hit in a list, though it appeared lower down in the lists sometimes. Also, no reasonable explanation emerged for why the protein had strikingly different molecular masses in different clonal populations. To add to the confusion, mass spectrometry strangely failed to offer any sort of alternate protein ID for the Lc3b^No^ bands. (Mass spectrometry results are provided in Supplementary Tables **S1** and **S2**).

**Fig. 3:**
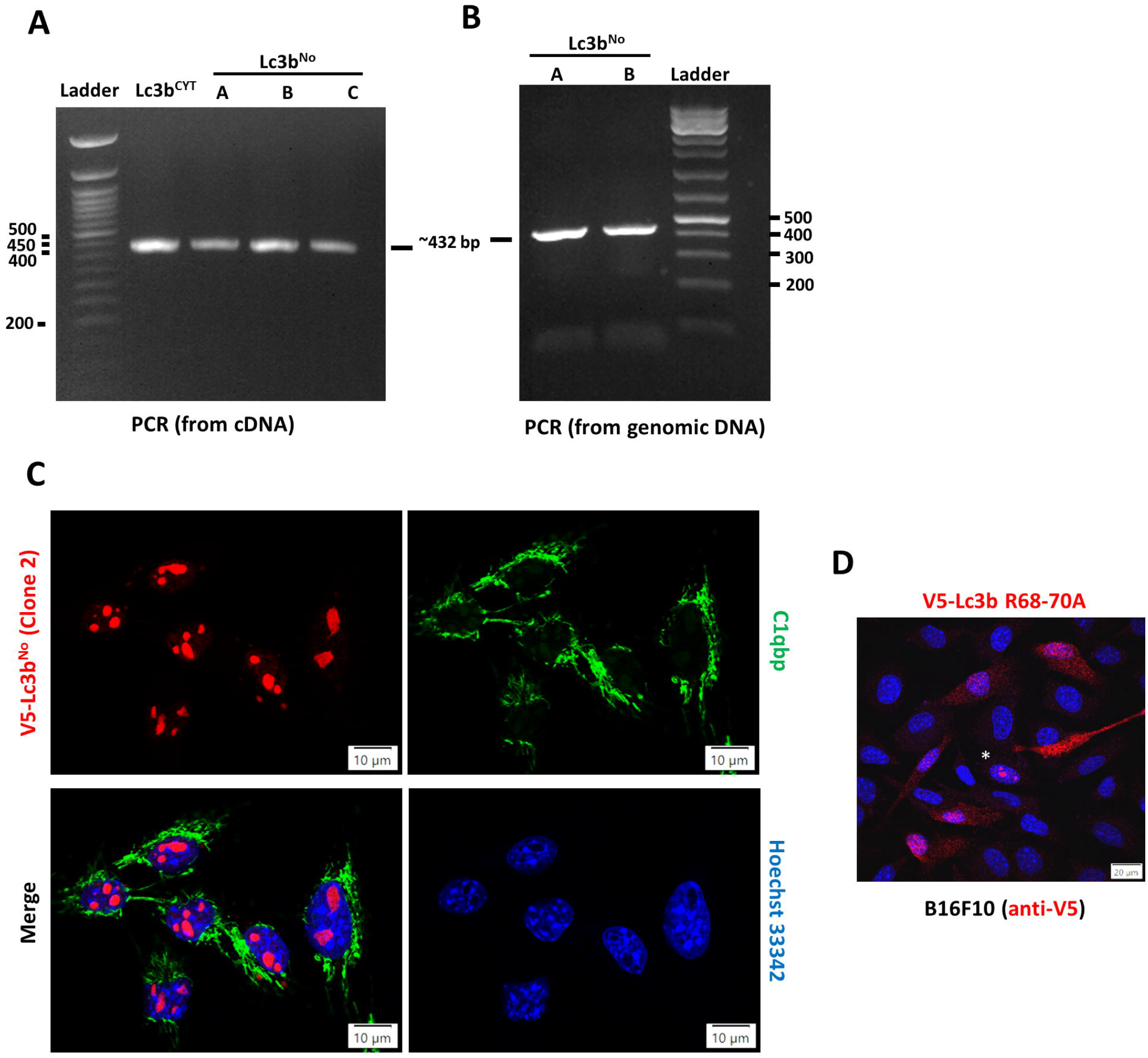
Data that urged us to re-think the conclusions made thus far. **A**. PCR with a V5-tag forward primer and Lc3b reverse primer on cDNA isolated from B16F10 cells stably expressing typical Lc3b (lane 1) or Lc3b^No^ (lanes 2-4). All reactions yielded a band corresponding to the size of the insert in the transfected construct. **B**. PCR with a V5-tag forward and Lc3b reverse primer with genomic DNA from B16F10 Lc3b^No^ clones 1 and 2 as template. The bands are similar to that in A. **C**. Double immunofluorescence of B16F10 cells stably expressing Lc3b^No^ with anti-V5 (red) and anti-C1qbp (green) antibodies. Nuclei are stained blue with Hoechst 33342. **D**. B16F10 cells transiently lipofected with an *Lc3b* construct in which residues R68-70 are mutated to alanines. A sporadic cell with nucleolar staining is asterisked.

Moreover, immunofluorescence staining in the Lc3b^No^ clones detected C1qbp primarily in the mitochondria and V5-Lc3b^No^ in the nucleoli **(Fig. 3C)**, presenting a conundrum as to how, and why, would two proteins spatially separated in compartments within the cell interact so strongly. There was weak staining of C1qbp in the nucleoli (you need to squint to see it in Fig. 3C), but the quantity was by far eclipsed by that of the protein in the mitochondrial matrix. It appeared more likely that the two proteins were interacting post cell lysis, during which they would be dislodged from their respective compartments. Kraft et al. had earlier reported that the mutation of a triple arginine motif in Lc3b resulted in complete loss of nucleolar localization (Kraft et al., 2016). But, in our case, even a R68-70A-mutated *V5-Lc3b* showed the typical number of sporadic cells with nucleolar localization post transfection on a cover slip **(Fig. 3D)**, suggesting something amiss.

Further, though this phenomenon was consistently reproducible by cloning *V5-Lc3b* from scratch, we found that several rounds of screening cell lines stably expressing *Lc3b* tagged with other small epitope tags – HA-tag, myc-tag, or a 3xFLAG tag – failed to produce a clone with nucleolar LC3b.

These were red flags; thus we looked at alternate explanations for this phenomenon.

### 3.4 The protein targeted to the nucleoli is a chimera comprising parts of Lc3b and a frameshifted sequence of puromycin N-acetyltransferase

Until this point, we had considered hypotheses pertaining to post-translational modification or epigenetic modification of genome-integrated Lc3b leading to its nucleolar localization. But now we concluded that these were unlikely, due to lack of any sort of supporting evidence and the contradictions mentioned above. Collating the available data, we decided that the next most likely hypothesis was that a chimeric protein comprising some part of Lc3b might be responsible for this phenomenon. If so, the correct-size PCR products of *V5-Lc3b* from the nucleolar clones were probably reflective of multiple integrations of the construct in the genome post transfection. At least one Lc3b^No^ clone also weakly stained for V5-Lc3b in some ALIS-like structures in the cytoplasm, suggesting possible expression from multiple loci. This is also reflected in the additional bands at the position where typical Lc3b would be expected to run, in the blot lanes for Lc3b^No^ clone 2 in Fig. 2 A, B and E.

If indeed a hypothetical chimeric protein was what was responsible for the nucleolar localization, it obviously had the V5-tag at the N-terminus, but the C-terminus of the chimeric sequence would be unknown and, therefore, not detectable as output in PCR-based experiments involving an Lc3b reverse primer. But the protein would still be detected in western blots. Since the molecular mass of LC3b^No^ from clone A was ∼20 kDa in blots (higher in other clones), the protein was longer than typical nonlipidated Lc3 (∼16 kDa) because it likely had a different C-terminus.

It was somewhat fortuitously that we managed to determine what constituted the C-terminus of the protein in nucleolar clone A.

Since the protein C1qbp interacted very strongly with the nucleolar protein and was showing up lower down the list in some MS/MS results from the 1D SDS-PAGE gels, we decided to check if V5-Lc3b^No^ was a chimera formed by illicit recombination between Lc3b and some part of C1qbp. C1qbp is known to exist as a trimer in cells (Jiang et al., 1999); this would also explain the interaction in the IP experiments. To this end, we used oligonucleotide primers at different positions along the sequence of the *C1qbp* gene (these primers were originally designed for a separate experiment seeking to determine which domain(s) of C1qbp was interacting with the nucleolar protein). We performed PCR with a V5-tag forward primer and each of these C1qbp primers, on cDNA from Lc3b^No^ clones A and B **(Fig. 4A)**. Curiously, a reverse primer coding for a region near one of the alpha helices of C1qbp (75-94) yielded a moderately strong band when clone A cDNA was used as template, though the size was shorter than what would be expected for that length of the C1qbp sequence. This band was excised from the gel and used to construct a clone in pBApo-EF1α Pur, by two-way overlap extension PCR with the forward primer of V5-Lc3b and the reverse primer of C1qbp, and subject to DNA sequencing (schematic in **Fig. 4B**). DNA sequencing revealed that the N-terminus of the protein comprised the V5-tag and nine N-terminal amino acids of Lc3b (MPSEKTFKQ), and the subsequent part of the fragment in the first overlapping product was, contrary to our expectations, neither from C1qbp nor from Lc3b but from an unknown sequence **(Fig. 4C)**. A deeper search revealed that it was part of a shifted reading frame of puromycin N-acetyltransferase (*pac*), starting from amino acid no. 56 **(Fig. 4E)**. It produced a match on an NCBI protein BLAST query, solely because an erroneous GenBank entry from the year 1999 (AAF01142.1; Gardner and Whitman, 1999) had the true N- and C-termini of puromycin N-acetyltransferase, but with a frameshifted region in the middle that corresponded to part of the ORF in this chimeric sequence. The correct complete sequence of puromycin N-acetyltransferase from *Streptomyces alboniger* **(Fig. 4D)**, first deduced by Lacalle et al. (Lacalle et al., 1989), is available from several other GenBank entries, such as the NCBI Reference Sequence WP_055528321.1.

**Fig. 4:**
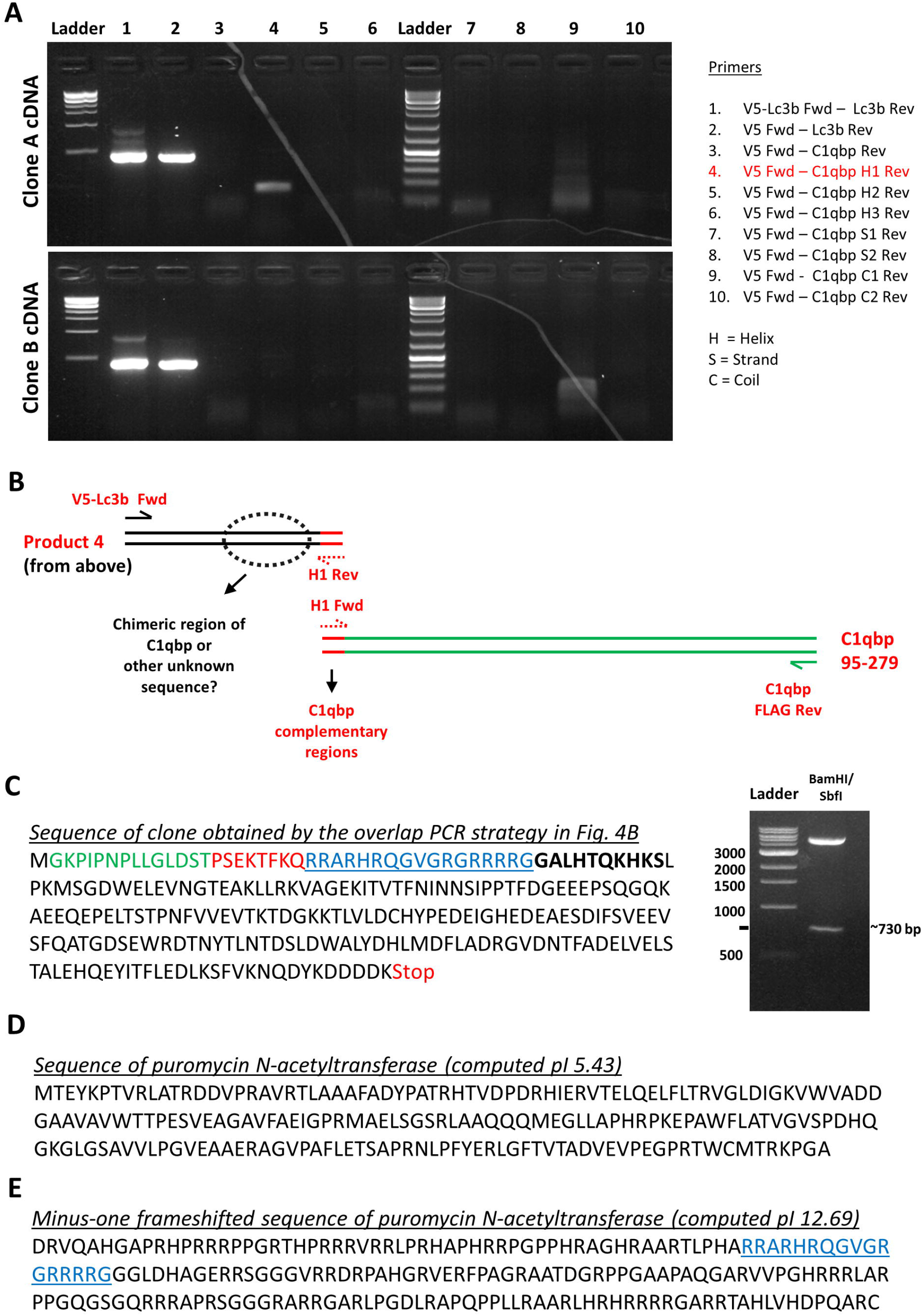
Figuring out the actual sequence of ‘Lc3b^No^’. **A**. PCR using cDNA from B16F10 Lc3b^No^ clones 1 and 2 as template, with the sets of primers mentioned on the right. Briefly, a V5-tag forward primer and different reverse primers along the length of the *C1qbp* gene were used to detect the possibility of an unlikely chimera. The primers were designed for a separate experiment to delete C1qbp helices, strands, and coil regions. The primer combination in lane 4 yielded a PCR product of ∼ 150 bp. The lowest band in the ladder on the left is 500 bp; the one on the right is a 100 bp ladder, with the brightest band 500 bp. **B**. Schematic of overlap extension PCR with complementary middle primers (H1 Fwd/H1 Rev) flanking C1qbp helix residues 75-94, forward primer of *V5-Lc3b*, and reverse primer of *C1qbp-FLAG*, with product from A as one of the two overlapping templates. **C**. Sequence of clone obtained through the strategy in B, with clone confirmation gel on the right (BamHI/SbfI). The amino acids of the V5 tag are indicated in green, Lc3b in red, a sequence unrelated to Lc3b in blue, and the parts of the C1qbp sequence in black. The ten black amino acids in bold indicate the position of the complementary primers flanking a helix deletion (amino acids 75-94) of C1qbp. **D**. Protein sequence of puromycin N-acetyltransferase from *Streptomyces alboniger* E. Minus one frameshifted sequence of puromycin N-acetyltransferase; amino acids in blue starting from no. 56 match the sequence from the clone in C.

### 3.5 Illegitimate recombination mediated by short homologies is the most plausible explanation for the formation of the chimeric sequence

Having discovered that the *Lc3b* and the *pac* sequences form a chimera, we knew that the *pac* sequence came from the drug resistance marker gene in the very plasmid which carried the *V5-Lc3b* insert – pBApo-EF1α Pur **(Fig. 5B)**. Using the *pac* sequence in the plasmid to follow the frameshifted sequence from its starting point in the chimera, the first stop codon was 148 amino acids downstream. This, combined with the V5 tag and the Lc3b N-terminus, would code for a protein of 171 amino acids, i.e. a molecular mass of ∼19 kDa – approximately the size of the V5-Lc3b^No^ from clone A.

**Fig. 5:**
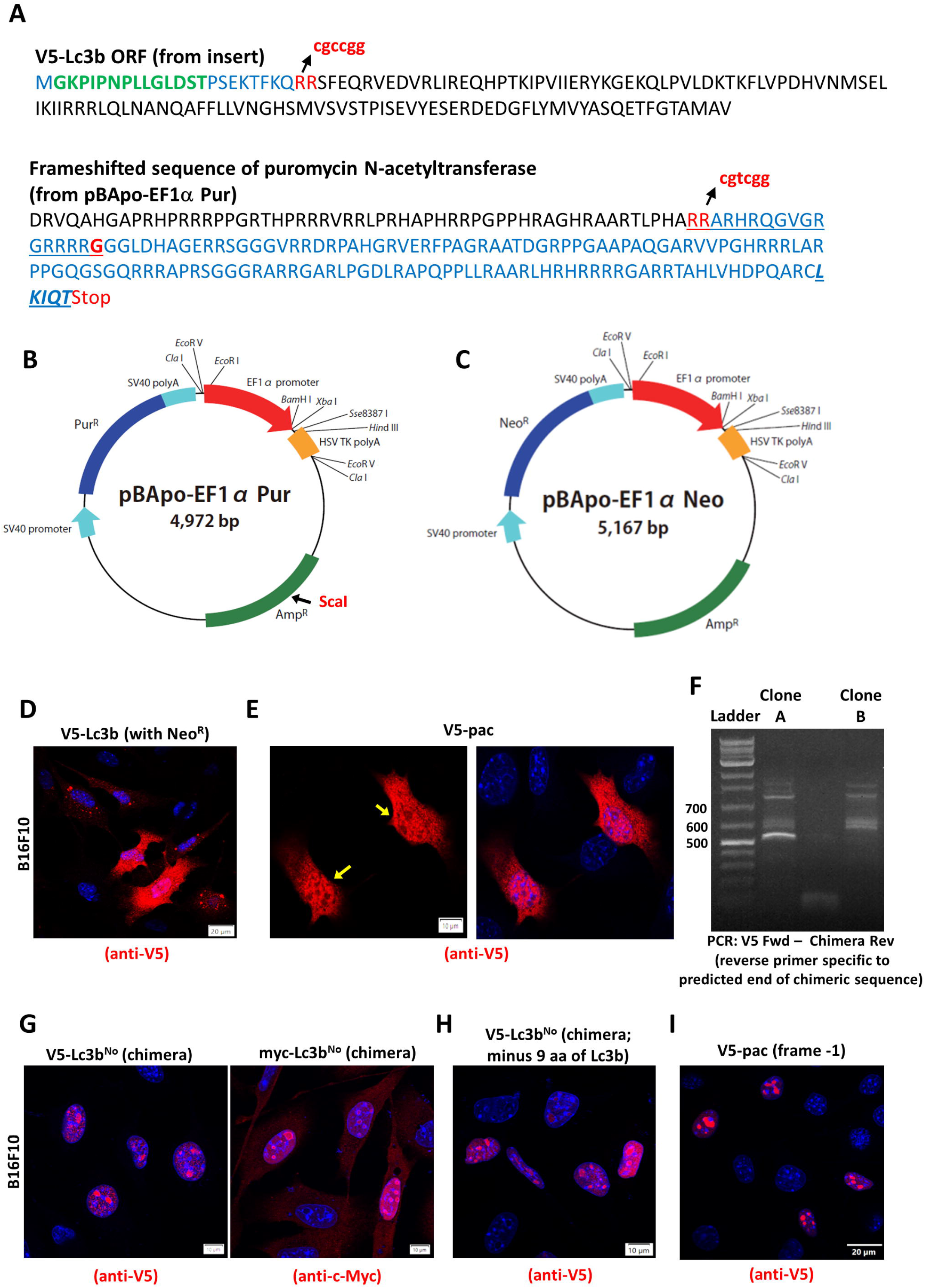
The frameshifted *pac* sequence is solely responsible for nucleolar targeting. **A**. Depiction of the sequences of the ORF of Lc3b and the frameshifted pac sequence (from pBApo-EF1α Pur) highlighting the parts of the final chimeric sequence in blue. The imperfect short homologies near the chimera junction points are shown in red. The bold G indicates the point where the C1qbp H1 reverse primer primed the reaction in Fig. 4A, enabling the chimeric sequence to be detected. The italicized and underlined blue amino acids at the end of the chimeric sequence are derived from the plasmid pBApo-EF1α Pur. **B**. Diagram of the plasmid pBApo-EF1α Pur (from the Takara product manual), showing important features. The ScaI site used for linearization prior to transfection is indicated in red. **C**. The plasmid pBApo-EF1α Neo, with a Neo^R^ instead of Pur^R^. All the remaining constructs in this figure were cloned in this plasmid. **D**. The *V5-Lc3b* construct cloned in pBApo-EF1α Neo transiently lipofected into B16F10 cells and detected with anti-V5 antibody; a representative field. No cells with nucleolar staining were found in any field. **E**. The correct ORF of *pac* cloned in pBApo-EF1α Neo and lipofected into B16F10 cells. pac shows a cytoplasmic and nuclear staining pattern, while being excluded from the nucleolus (yellow arrows). **F**. PCR on genomic DNA from B16F10 Lc3b^No^ clones 1 and 2 with a *V5*-tag forward primer and a reverse primer complimentary to the vector-derived end of the chimeric sequence. **G**. Exact sequence of the chimera from clone 1, cloned in pBApo-EF1α Neo (with two different epitope tags) and lipofected into B16F10 cells. **H**. The chimeric sequence lacking the nine amino acids of Lc3b cloned into pBApo-EF1α Neo and lipofected into B16F10 cells. **I**. The full -1 frameshifted sequence of *pac* was cloned into pBApo-EF1α Neo and lipofected into B16F10 cells.

Next, we investigated how the two reading frames could come together. *V5-Lc3b* had been cloned by precisely excising a ∼430 bp PCR product from a gel; therefore, contamination from a higher molecular mass cDNA-derived insert, or any other cloning artefact, is ruled out. In pBApo-EF1α Pur, the sequence between the BamHI site used to clone *V5-Lc3b* and the start of the chimeric region (in the *pac* gene) is >3000 bp long, and illegitimate recombination resulting in deletion of the intervening sequence during/after integration of the plasmid in the host genome is one plausible explanation for this phenomenon. Also, homogeneous clonal populations usually reflect a heritable change in the DNA. On analysing the chimera junction point (recombination site) in both sequences, it was noted that the same two amino acids – RR – are encoded by the nucleotide sequence at both the sites, but with imperfect repeats – the nucleotide sequence in *Lc3b* being 5’-cgccgg-3’, and the one in the *pac* reading frame being 5’-cgtcgg-3’ **(Fig. 5A)**. Nonhomologous recombination involving short homologous repeats and deletions of the intervening sequences, at the end of which only one of the repeat sequences is retained in the parent molecule, is a well characterized phenomenon in prokaryotes and eukaryotes (Franklin, 1967; Bi and Liu, 1996; Brunier et al., 1989; Bullock et al., 1985). In this case, the sequence of -RR- in the final recombined sequence read 5’-cgccgg-3’, suggesting that the residues from the *Lc3b* sequence were retained in the chimeric product. An alternate means for this event to occur would be crossover recombination between two or more molecules of the transfected plasmid substrate at these regions of short homologies during integration into the genome.

It is worth noting that we were able to observe nucleolar localization of the V5-tagged protein both when linearized plasmid was transfected into cells for stable expression (in clones after drug selection), and also when circular plasmid was transfected into cells for transient expression (in isolated cells), indicating that linearization of the substrate prior to transfection was not a prerequisite for this event to occur in cells. Since the restriction enzyme site used for plasmid linearization was located midway between the multiple cloning site and the puromycin marker gene, **(Fig. 5B)** recombination could have proceeded through one or more of the mechanisms discussed above, depending on whether the plasmid was linearized or circular, with the end result being the formation of chimeric sequences at different regions having short homologies. Indeed, illegitimate recombination between short homologies can occur by a variety of sophisticated mechanisms in mammalian cells (van Rijk and Bloemendal, 2003), the discussion of which would be beyond the scope of this study.

To understand things better, we cloned *V5-Lc3b* in pBApo-EF1α Neo, the sister plasmid of the one originally used **(Fig. 5C)**. This contains a neomycin resistance marker instead of *pac*. Like we suspected, not a single cell stained for V5-Lc3b in the nucleoli post-transfection, indicating that the *pac* gene was integral to the manifestation of this phenomenon **(Fig. 5D)**. In the same plasmid pBApo-EF1α Neo, we cloned the complete correct ORF of *pac* with a V5 tag at the N-terminus. Immunofluorescence staining showed that the pac protein was majorly cytosolic and nuclear, but excluded from the nucleoli **(Fig. 5E)**, confirming that the puromycin N-acetyltransferase protein itself has no tendency to localize to the nucleolus, but a -1 shift in the reading frame of the same sequence is needed to confer upon the protein that property.

Next, we attempted to reproduce the exact phenomenon in the most direct manner, by cloning the full sequence of the nucleolus-localized chimera in pBapo-EF1α Neo — a V5 tag at the N-terminus, and nine amino acids of Lc3b followed by the frameshifted *pac* sequence (amino acids 56 -198; 143 amino acids from the *pac* sequence), and another similar construct with a myc-tag instead of V5 at the N-terminus. For this, we used an Lc3b forward primer containing the respective tag, and a reverse primer at the predicted end of the chimeric sequence. PCR from genomic DNA of clone A yielded a clear band of the predicted amplicon **(Fig. 5F)**, confirming the presence of the chimeric DNA in the genomic DNA of clone A. Both V5- and myc-tagged constructs produced proteins intensely staining the nucleoli when transiently transfected into B16F10 cells **(Fig. 5G)**. This confirmed two things – that the chimeric ORF was responsible for nucleolar targeting and that the epitope tag had no role in nucleolar targeting. Finally, we made another construct with only the V5-tag and the shifted reading frame of *pac*, but without the N-terminal nine residues of Lc3b. This, too, showed the same pattern **(Fig. 5H)**, indicating that the nine residues of Lc3b had no function in nucleolar targeting either, and that this was mediated entirely by the frameshifted protein sequence from *pac*.

The full *pac* ORF codes for a 199-amino acid protein. Curiously, the minus-one shifted frame of the gene, though not an ORF, also codes for as many uninterrupted amino acids, with no out-of-frame (‘ambush’) stop codons within this reading frame. This might be partly due to the fact that *Streptomyces* genomes are GC-rich and stop codons are AT-rich. However, none of the other antibiotic resistance genes commonly used in plasmids have entire frameshifted sequences uninterrupted by stop codons. Since the chimera in this study only coded for the latter ∼143 amino acids of the frameshifted sequence, we cloned the entire 199 amino acid minus-one frameshifted *pac* (with an added methionine at the start) in pBApo-EF1α Neo. We found that there was no change in localization of this full-length frameshifted product compared with the chimera, and that it was heavily enriched in the nucleoli **(Fig.5I)**.

## Discussion

Map1Lc3b is a protein that never ceases to be discussed. Its primary role in autophagy thrust it into perpetual limelight, and it was always known to be present in both the nucleus and the cytoplasm of cells. But its function in the nucleus only became apparent rather recently, when Huang et al. reported in 2015 that it was activated in the nucleus by the deacetylase Sirt1 in order to redistribute to the cytoplasm and take part in its important functions in autophagy (Huang et al., 2015). Reports of its presence in the nucleolus arrived only later. In 2012, when we first obtained these clones with the protein abundantly enriched in the nucleoli it seemed to us like new, then later corroborative, data.

But the functions of Lc3b in the nucleoli of cells, if any, still remain elusive. You might recall from Fig.1G that Lc3b tagged with GFP and mCherry transiently transfected into cells did display some staining in the nucleoli in addition to other parts of the cell, but the fact that this is not the case when smaller tags are used points to the fluorescent fusion proteins interfering with either the distribution or normal diffusion kinetics of Lc3b.

In this study, the ‘nucleolar localization’ of Lc3b observed by us turned out to be different from those reported elsewhere. Despite being detected by anti-Lc3 antibodies, it was a chimeric protein resulting from illegitimate recombination between a portion of Lc3b and a shifted (−1) reading frame of puromycin N-acetyltransferase that was being targeted to the nucleolus, with the property of nucleolar targeting imparted by the latter. One important thing to note here is that we determined the chimeric composition of the protein in only one of our clones (∼20 kDa). We were fortunate that the use of a totally unrelated oligonucleotide primer made it possible to capture the sequence of the chimera. It is likely that the other clones where the molecular mass of the nucleolar protein is ∼37 kDa and above have different chimera junction points and comprise shorter or longer lengths of either Lc3b or/and the pac sequence, or possibly multiple repeats of the sequences (due to their larger size). In the ∼20 kDa clone, the chimera junction point corresponded to codons for a double arginine in both sequences. One look at the frameshifted pac protein sequence (Fig. 4E) reveals that there are multiple stretches of two or more arginines (thirteen, to be exact). Recombination at any of these residues and/or at different segments of the *Lc3b* ORF coding for other amino acids might have produced chimeras of different lengths in the other clones, leading to a similar end-result (targeting to nucleoli), which we decided was not worth investigating because it was unlikely to give us any additional insight of value.

### The -1 frameshift massively alters the pI of the pac protein

The sheer number of arginine stretches in the amino acid sequence of the frameshifted pac is indicative of the protein likely to have affinity to the nuclear/nucleolar compartments (though the protein is originally from an organism that does not possess a nucleus). Martin et al. have reported that peptide entities consisting of only arginines, or multiple arginines accompanied by an NLS, with an isoelectric point (pI) of 12.6 or above are sufficient to mediate nucleolar accumulation (Martin et al., 2015). Intriguingly, we find that a minus-one frameshift of the 199-amino acid *pac* sequence drastically alters the theoretically computed pI (Expasy Compute pI/Mw Tool ; Duvaud et al., 2021) of the translated protein from 5.43 to 12.69, which falls perfectly at the requisite pI threshold mentioned for charge-dependent nucleolar accumulation.

C1qbp/p32 is a protein that is primarily present in the mitochondrial matrix but has been shown to have a presence, and important functions, in the nucleolar compartment. C1qbp is a multifunctional protein with a large interactome and has, in at least two other reports, been shown to interact with an arginine stretch in a partner protein (Xu et al., 1999; Wang et al. 2019). Thus, it is likely that the very strong interaction of the C1qbp with the protein in this study is mediated by interaction between the acidic residues of C1qbp and one or more stretches of arginine residues in the frameshifted pac sequence. The robustness of this interaction, coupled with the fact that the frameshifted protein of *pac* is not a protein that naturally occurs in any living systems, makes the latter a potential candidate for use in affinity pulldown experiments that use C1qbp as a bait protein. Another potential application of the frameshifted pac protein would be to deploy it (or even a partial stretch of the protein) as a fusion partner if enrichment or sequestration of a target protein in the nucleolus is desired for a specific application.

### The V5 tag signifies the importance of context in nonhomologous recombination events

It has been reported previously that deletion termini in eukaryotes have sequence features that are less striking than those in prokaryotes, with repeats as few as 2 to 3 base pairs in length contributing to spontaneous deletions (Nalbantoglu et al., 1986; Roth and Wilson, 1986; Ruley and Fried, 1983, Pulak and Anderson, 1988), whereas significantly more homology is required in prokaryotes. However, it also appears that not only short homologous sequences but also the context of the surrounding regions in the DNA are hugely important in ‘nonhomologous’ illegitimate recombination in eukaryotes. Despite screening hundreds of clones, the presence of small epitope tags other than the V5 tag at the N-terminus of Lc3b did not yield any cells or clonal populations expressing the tagged protein in the nucleolus. It is important to make the distinction that the transfection of a construct encoding a *tag*-*Lc3b*-*pac* chimeric ORF resulted in the expressed protein localizing in the nucleoli regardless of the type of epitope tag, but de novo recombination of an *Lc3b* construct with the *pac* sequence to yield a chimeric ORF happened only when the transfected *Lc3b* construct had a V5 tag at the N-terminus. The V5 tag seems to assist this unlikely recombination event in some unknown manner and increases the probability of its occurrence to the levels reported earlier in this text.

### What is the general likelihood that this type of phenomenon will occur in routine molecular biology experiments involving DNA and transfections?

Phenomena such as the one reported in this paper stress the need for scientists to be careful when interpreting unusual or stand-out results from experiments, because all sorts of artefacts abound in biological systems – and especially when molecular biology tools are used to investigate or mediate in cellular phenomena.

We have used the plasmid pBApo-EF1α Pur to express many genes over the years, but so far have not noticed an aberration similar to this in any other experiment. Hundreds of plasmid vectors used by scientists across the world contain the puromycin resistance gene as a selection marker and, by implication, the reading frame of the nucleolus-targeted protein. Therefore, it would not be a surprise if this exact type of mislocalization manifests itself in some other experimental context, with some other gene construct, provided all favourable factors come together as they did here. Perhaps it is yet another sign that experiments involving random and unquantifiable integrations of DNA in the genome should gradually become a thing of the past, as more and more targeted methodologies become available for gene expression and modification that promise specificity and reproducibility. At the same time, the very low threshold that has been reported for nonhomologous recombination-based deletions to occur in eukaryotes, with very little homology required as demonstrated in this study, calls for the need to be extremely vigilant during gene editing and other applications where DNA modification is the primary objective. Inevitably, applications such as DNA vaccination also come to mind. The necessity to screen thoroughly for potential off-target effects can never be emphasised enough, especially because not all artefactual phenomena provide a visible output.

If homology at as few as one or two codons suffice for recombination to be triggered in eukaryotes, characterising the accessory cis-acting factors (akin to the V5 tag in this study) that promote/prevent recombination events become very important because there are innumerable loci that would be candidates for recombination but most of them do not engage in it, ensuring that the sanctity of the genome remains mostly intact, maintaining order amidst chaos. Also, such phenomena and the varying mathematical probabilities associated with them lead us to reflect about how frequently or infrequently such events occur in different cells and tissues, and how they might be shaping the evolution of genomes in several small steps over large time frames.

## Materials and Methods

### Cell lines and reagents

B16F10 mouse melanoma was obtained from the tissue culture facility, Dabur Research Foundation, Ghaziabad, India. PC3 prostate cancer cells (ATCC® CRL-1435) and BeWo choriocarcinoma cells (ATCC® CCL-98) were gifts from Prof. Sathees C Raghavan and Prof. R Manjunath, Indian Institute of Science, Bangalore, India. B16F10 melanoma was cultured in high glucose Dulbecco’s Modified Eagle’s Medium (DMEM) containing 10% foetal bovine serum (FBS) and an antibiotic mixture comprising 10,000 units mL^-1^ penicillin, 10,000 μg mL^-1^ streptomycin, and 25 μg mL^-1^ amphotericin B. PC3 cells were cultured in Kaighn’s modification of Ham’s F12 medium containing the same additives as DMEM. The cell lines were maintained in an incubator at 37°C with 5% CO_2_ and water provided for humidification. All cell lines were routinely inspected for the absence of any potential contamination.

DMEM, FBS (E.U.-approved, South America origin), Earle’s Balanced Salt Solution (EBSS), Opti-MEM Reduced Serum medium, Antibiotic-Antimycotic solution (100X), Pierce BCA protein assay kit, PageRuler prestained protein ladder, GeneJET plasmid miniprep and gel extraction kits, and ProLong™ Glass antifade mountant were purchased from Thermo Fisher Scientific, Waltham, Massachusetts, USA. Trypsin (from porcine pancreas), puromycin dihydrochloride, Lipofectamine 2000, TRI reagent for RNA isolation, and Hoechst 33342 were bought from Sigma-Aldrich, St. Louis, MO, USA. Glutathione resin was purchased from G-Biosciences, St. Louis, Missouri. Bovine albumin, fraction V, fatty acid-free was purchased from MP Biomedicals Ltd., Navi Mumbai, Maharashtra, India. All oligonucleotides used in PCR and cloning were procured from Sigma-Aldrich. Phusion® polymerase, DNA ladders and all restriction endonucleases used in cloning protocols were procured from New England Biolabs (NEB), Ipswich, Massachusetts, USA. Taq DNA polymerase was from Genei Labs, Bengaluru, India. cOmplete™ protease inhibitor cocktail tablets, EDTA-free, were purchased from Roche, Basel, Switzerland. Coomassie Brilliant Blue R250 and high purity acrylamide were purchased from Sisco Research Laboratories, Mumbai, India. Seakem® LE Agarose was from Lonza, Rockland, ME, USA. Sanger DNA sequencing was performed at Eurofins Scientific, Bengaluru, India.

### Antibodies

The following antibodies were used in this study: anti-V5 (mouse monoclonal, R960-25, Thermo Fisher Scientific), anti-FLAG (mouse monoclonal, F1804, clone M2, Sigma-Aldrich), anti-LC3B L7543, rabbit polyclonal, Sigma-Aldrich), anti-C1qbp (PAA651Mu01/Hu01, Cloud-Clone Corp., TX, USA) anti-β-actin (A3854, rabbit polyclonal, HRP-conjugate, Sigma-Aldrich), anti-V5 agarose affinity gel (A7345, monoclonal clone V5-10, Sigma-Aldrich), anti-Fibrillarin (ab5821, Abcam, MA, USA), anti-Nucleophosmin (FC82291, Abcam, MA, USA), anti-c-Myc (OP10, EMD Millipore Corp., MA, USA) and anti-HA (S3143, Epitomics, CA, USA).

### Plasmids, constructs, and sequences

The *Map1Lc3b (Lc3b)* gene was amplified by PCR from cDNA of NIH 3T3 mouse fibroblast cells. The *V5-Lc3b* construct was made by cloning the insert into the BamHI and HindIII sites of the vector pBApo-EF1α Pur, and subsequently also in pBApo-EF1α Neo (Takara Bio Inc., Kusatsu, Shiga Prefecture, Japan). The V5 sequence was included in the forward primer. *myc*-*Lc3b* and *HA*-*Lc3b* were constructed likewise in the same vector. *3xFLAG-Lc3b* was made by cloning the *Lc3b* sequence into the HindIII and KpnI sites of the vector p3xFLAG-CMV-10 (Sigma Aldrich). *mCherry-Lc3b* and *GFP-Lc3b* were made by cloning *Lc3b* into the EcoRI and BamHI sites of pEGFP-C1 (Clontech Laboratories, CA, USA). GST-Lc3b was constructed by cloning *Lc3b* into the BamHI and SalI sites of pGEX-4T-1 (GE Healthcare, IL, USA). The pac sequence was amplified from pBApo-EF1α Pur and cloned into pBApo-EF1α Neo. All constructs described in Section 5 were cloned in pBApo-EF1α Neo into its BamHI and HindIII sites. All mutations were created by an overlap extension PCR strategy using oligonucleotide primers containing the desired mutations.

DNA sequencing results were analysed using the Translate tool at the Expasy portal (Swiss Institute of Bioinformatics (Lausanne, Switzerland).

### Creation of cell lines stably expressing proteins

Briefly, the *Lc3b* construct in pBApo-EF1α Pur was linearized using ScaI and transfected into semi-confluent B16F10 cells using Lipofectamine 2000. 48 h later, the cells were subcultured into six-well plates and transformants were selected with 2 μg mL^-1^ puromycin (B16F10 cells) or 1 μg mL^-1^ (PC3 cells). 10-14 days later, single colonies were picked using sterile PYREX® cloning cylinders (6 mm x 8 mm, Corning Inc., Corning, New York, USA) and expanded into cultures in 6-well plates. The colonies were screened for Lc3b expression by indirect immunofluorescence using anti-V5 antibody, and clones with the desired expression patterns/levels were further expanded and frozen. Likewise, to screen for presence of nucleolar localization, the same procedure was performed with the various constructs mentioned, except that transfected cultures were plated on 6-well plates containing cover slips in the wells. After immunofluorescence staining, the cover slip was systematically scanned under the microscope to search for colonies exhibiting nucleolar staining. In this case, the colonies were imaged but not isolated or passaged.

### Transfection procedures

Lipofectamine 2000 was used according to the manufacturer’s instructions. Opti-MEM Reduced Serum medium was replaced with fresh medium 4-6 h post electroporation, and the cells were processed for downstream experiments 16-24 h post transfection for transient procedures, or cultured for longer periods under drug selection to create cell lines stably expressing the constructs.

### Immunoprecipitation and pull-down

For immunoprecipitation of cells lysed under denaturing conditions, cells in P100 plates were washed with PBS, scraped into tubes with the help of PBS, and centrifuged at 400 *g* for 10 minutes. They were lysed with denaturing lysis buffer containing 50 mM Tris.Cl pH 7.4, 5 mM EDTA, 1% (w/v) SDS, and 10 mM DTT (100 μL of buffer per 0.5-2 × 10^7^ cells). After vortexing and boiling for 5 min., the denatured protein extract was diluted ten-fold with nondenaturing lysis buffer containing 50 mM Tris.Cl pH 7.4, 300 mM NaCl, 5 mM EDTA, 1% (w/v) Triton X-100, and protease inhibitors, followed by sonication to shear DNA. The lysate was centrifuged at 10,000 rpm for 15 minutes at 4 C and incubated with 20-30 μL of anti-V5 agarose affinity gel (or anti-C1qbp, followed by Protein A agarose beads) to immunoprecipitate the target protein. After incubation for 2 h, the beads were washed thrice with wash buffer and were ready for SDS-PAGE.

GST pull-down assays, SDS-PAGE, and western blotting were performed essentially as described elsewhere (Thekkinghat et al., 2019).

### Indirect immunofluorescence

Cells grown on acid-washed coverslips were fixed with 3.7% formaldehyde, washed twice with PBS, and permeabilized with 0.1% Triton X-100 in PBS. After blocking with 5% BSA in PBS for 30 min., they were incubated with primary antibodies in blocking solution for 2 hours. Most antibodies were used at dilutions between 1:200 to 1:500. After 3 washes with PBS containing 0.05% Triton X-100, they were incubated with secondary fluorophore-conjugated antibodies (Alexa Fluor 555/488; Thermo Fisher Scientific/CST; 1:200 dilution) in blocking solution for 2 hours. After 3 more washes, they were stained with 0.2 μg mL^-1^ Hoechst 33342, washed once, and mounted on a drop of ProLong® Glass antifade mountant.

Imaging was done on an Olympus Fluoview FV3000 confocal laser scanning microscope. Image processing was done using the associated Olympus cellSens Dimension software. Cropping, arranging and labelling of images were done using a combination of Microsoft PowerPoint and GNU Image Manipulation Program (GIMP).

### Mass spectrometry

Protein bands stained with Coomassie Brilliant Blue were excised precisely from SDS polyacrylamide gels and subjected to trypsin digestion and mass spectrometry, raw data analysis and processing using MaxQuant software (v1.6.2.10) and MS Excel essentially as described elsewhere (Tushir et al., 2021), but against the *Mus musculus* proteome.

## Acknowledgements

The first author (A. A. T) is indebted to his PhD supervisor, P. N. R for graciously permitting him to work in the laboratory and use the laboratory resources for this project, both during and after his PhD, as well as for helpful suggestions, uninhibited freedom to experiment, and constant encouragement. The services of the confocal microscopy facility, Dept. of Biochemistry, Indian Institute of Science, and the operators Ms Puja Biswas and Ms Spoorthi are acknowledged. For mass spectrometry services, the authors are grateful to Prof. Utpal Tatu and Sathisha Kamanna, Dept. of Biochemistry, IISc and the staff at Technology Platform Services, C-CAMP, Bangalore. The authors also thank Dr. Pritha Mukherjee for her enthusiastic contribution in cloning the *V5-Lc3b* construct when she was a summer trainee in the lab many years ago. A. A. T is grateful for the unconditional friendship and support of all the members of the Rangarajan lab.

## Author Contributions

A.A.T and P. N. R conceptualized the study. A. A. T performed the experiments, validated and analysed the results, verified the reproducibility of the data, wrote and edited the original draft of the manuscript. A. A. T and P. N. R reviewed and edited the manuscript. P. N. R provided resources, funding, and supervision.

## Data Availability Statement

The sequences of all oligonucleotide primers used in this study will be made available on request.

